# Novel insights into the mode of action of 1,4-dioxane using a systems screening approach

**DOI:** 10.1101/2020.12.27.424470

**Authors:** Georgia Charkoftaki, Jaya Prakash Golla, Alvaro Santos-Neto, David J. Orlicky, Rolando Garcia-Milian, Ying Chen, Nicholas J.W. Rattray, Yuping Cai, Yewei Wang, Colin T Shern, Varvara Mironova, Yensheng Wang, Caroline H. Johnson, David C. Thompson, Vasilis Vasiliou

**Author notes:** **Corresponding Author:** Vasilis Vasiliou, PhD, Department of Environmental Health Sciences, Yale School of Public Health, 60 College Street, Rm. 511, PO Box 208034, New Haven, CT 06520-8034. Current address: Department of Medicine, Yale University School of Medicine, New Haven, CT 06520, United States; Veterans Affairs Connecticut Healthcare System, West Haven, CT 06516, United States.

## Abstract

1,4-Dioxane (1,4-DX) is an environmental contaminant found in drinking water throughout the United States (US). While it is a suspected liver carcinogen, there is no federal or state maximum contaminant level for 1,4-DX in drinking water. Very little is known about the mechanisms by which this chemical elicits liver carcinogenicity. In the present study, female BDF-1 mice were exposed to 1,4-DX (0, 50, 500 and 5,000 mg/L) in their drinking water for one or four weeks, to explore the toxic effects. Histopathological studies and a multi-omics approach (transcriptomics and metabolomics) were performed to investigate potential mechanisms of toxicity. Immunohistochemical analysis of the liver revealed increased H2AXγ-positive hepatocytes (a marker of DNA double strand breaks), and an expansion of precholangiocytes (reflecting both DNA damage and repair mechanisms) after exposure. Liver transcriptomics revealed 1,4-DX-induced perturbations in signaling pathways predicted to impact the oxidative stress response, detoxification, and DNA damage. Liver, kidney, feces and urine metabolomic profiling revealed no effect of 1,4-DX exposure, and bile acid quantification in liver and feces similarly showed no effect of exposure. We speculate that the results may be reflective of DNA damage being counterbalanced by the repair response, with the net result being a null overall effect on the systemic biochemistry of the exposed mice. Our results show a novel approach for the investigation of environmental chemicals that do not elicit cell death but have activated the repair systems in response to 1,4-DX exposure.

## Introduction

1,4-Dioxane (1,4-DX) is a synthetic industrial chemical used as a stabilizer for chlorinated solvents and as a solvent in a variety of organic products, such as dyes, greases, varnishes, waxes, detergents, cosmetics, and deodorants (EPA, 2017). It has been inadvertently released into the environment during its production, the processing of other chemicals, and its use during the manufacture of consumer products. It migrates rapidly into the ground water and is resistant to hydrolysis by water and to microbial degradation. The most significant sources of 1,4-DX in drinking water are wastewater discharge, unintended spills and leaks, and historical disposal practices of its host solvent, e.g., trichloroacetic acid (TCA). 1,4-DX is also found as an impurity in some consumer products, such as deodorants, shampoos, and cosmetics (Wilbur *et al.*, 2012). Trace levels of 1,4-DX may be present in manufactured food additives, food (due to absorption from 1,4-DX-containing adhesives on packaging material), or on food crops treated with pesticides that contain 1,4-DX as a solvent (EPA, 2017).

Due to is physicochemical characteristics, 1,4-DX is resistant to degradation by natural or engineered processes. Its low organic phase and water partition coefficient makes it highly miscible in water, unlikely to retard or adsorb to a solid phase, and to have a low tendency to volatilize (Cashman *et al.*, 2019). These properties are unfavorable for conventional treatment strategies (Favara *et al.*, 2016; Stepien *et al.*, 2014). Therefore, 1,4-DX can leach into water systems and accumulate in groundwater sites. Its low mobility in clay soils contribute to a long half-life of 2-5 years in groundwater (Khan *et al.*, 2018; Pollitt *et al.*, 2019). Reflective of these issues, a recent study showed that 1,4-DX can be detected in 21 % of public drinking water (Adamson *et al.*, 2017).

Several studies have shown that 1,4-DX can cause hepatic toxicity and carcinogenicity in mice and humans (Carrera *et al.*, 2017; Kasai *et al.*, 2009; Kraybill, 1978). Single cell necrosis and swelling of centrilobular hepatocytes was observed in male and female Crj:BDF-1 mice exposed to ≥ 4,000 ppm 1,4-DX in drinking water for 13 weeks (Kano *et al.*, 2008). After exposure to a lower dose (1,600 ppm) of 1,4-DX for the same period of time, only females exhibited increased nuclear enlargement in the bronchial epithelium. These results suggest a sex difference in sensitivity to the toxic effects of 1,4-DX. Sex-differences were also observed in 1,4-DX sensitivity in a longer term (2 year) study in Crj:BDF-1 mice. Hepatocellular carcinoma incidence increased in female mice at 500 and 2,000 ppm 1,4-DX in drinking water while the incidence increased only at 2,000 ppm in male mice (Kano *et al.*, 2009). Interestingly, the carcinoma incidence in mice of either sex was not increased (relative to control) after exposure to 8,000 ppm 1,4-DX in drinking water for 2 years (Kano, Umeda, Kasai, Sasaki, Matsumoto, Yamazaki, Nagano, Arito and Fukushima, 2009)

Given its apparent potential for causing liver damage, and for contaminating drinking water and food, 1,4-DX is a health concern. The long half-life of 1,4-DX in groundwater and lack of current remediation strategies makes it even more environmentally challenging. Although there is evidence linking 1,4-DX to liver cancer, the mechanisms by which it induces cancer are not known. In the present study, we exposed female BDF-1 mice to a range of 1,4-DX doses (0, 50, 500 and 5,000 ppm) in drinking water for one or four weeks, and used a combination of metabolomic, transcriptomic and histopathological analyses to delineate the hepatic and broader systemic biological effects of 1,4-DX in female BDF-1 mice.

## 2. Methods and materials

### 2.1 Chemicals

Cholic acid, chenodeoxycholic acid, hyodeoxycholic acid, sodium glycochenodeoxycholate, sodium taurochenodeoxycholate glycocholic acid hydrate, sodium taurolithocholic acid, sodium taurodeoxycholate hydrate, sodium taurohyodeoxycholate hydrate, and taurocholic acid sodium salt hydrate were purchased from Sigma-Aldrich (Saint Louis, MI, USA). Glycodeoxycholic acid sodium salt (EMD, Millipore Corp., Burlington, MA, USA), ursodeoxycholic acid, deoxycholic acid (ChemCruz, Santa Cruz Biotechnology Inc., Dallas TX, USA), lithocholic acid (Cayman Chemical Company, Ann Arbor, MI, USA), and glycodeoxycholic acid, glycolithocholic acid, glycodeoxycholic acid, 7-ketodeoxycholate and 7-ketolithocholic acid were purchased from Toronto Research Chemicals (North York, ON, Canada). Formic acid (99+%) was purchased in 1 mL ampules from Thermo Scientific (Rockford, IL). Ammonium formate and 2-propanol (both Optima^®^ LC/MS grade), acetonitrile and water (both Optima^®^ grade) were purchased from Fisher Chemical (Fair Lawn, NJ, USA).

### 2.2 Animal study

Three-week-old female BDF1 mice were purchased from Charles River (Wilmington, MA), housed in standard cages, and acclimatize for 7-10 days before initiation of the 1,4-DX exposure protocol. Mice were maintained on a 12 h light-dark cycle, with access to regular chow and water *ad libitum*. During the exposure protocol (one- or four-week duration), mice were provided with drinking water containing 1,4-DX (50, 500 or 5,000 mg/L) (n=6/group) or with drinking water alone (0 mg/L 1,4-DX, control) (n=4 mice). During these experiments, mice were housed two per cage. In order to evaluate the systemic effects of 1,4-DX exposure, measurements were taken of body weight, water consumption, exposure to 1,4-DX the liver transcriptome, and liver, kidney, urine and fecal metabolomes. Body weight was measured twice each week and the general condition of the mice was monitored continuously. Urine and feces samples were collected from mice individually placed in a metabolic cage (Nalgene, Techniplast Inc., Exton, PA) for 12 h. These samples were snap frozen in liquid nitrogen and stored at −80^°^C for latter metabolomic analyses. All mice were acclimatized to the metabolic cages at least twice before the actual sample collection. For mice exposed to 0 mg/L or 5,000 mg/L 1,4-DX for one week, feces were collected from four mice. Urine samples were collected from three control and five 5,000 mg/L 1,4-DX-exposed mice for one week, respectively. Feces samples were collected from three control and four 5,000 mg/L 1,4-DX-exposed mice; urine was collected from four mice in both groups. These differences in numbers were due to some mice not produced urine or feces during the 12 h period in the metabolic cage. After euthanasia by CO2 asphyxiation, the kidney and liver were rapidly harvested. A small piece of the liver and kidney was immediately fixed in 10% formalin for histologic evaluation and the remaining tissues were snap frozen using liquid nitrogen and stored at −80°C for latter metabolomic or transcriptomic analyses. All animal procedures were approved by and conducted in compliance with the Institutional Animal Care and Use Committee (IACUC) of Yale University.

### 2.3 Histopathology

Formalin-fixed, paraffin-embedded liver tissues from mice administered 0, 500, or 5,000 mg/L 1,4-DX for one week, and 0 or 5,000 mg/L 1,4-DX for four weeks were sectioned at 5 μm, and stained with hematoxylin and eosin (H&E), or subjected to immunohistochemistry (IHC) for the presence of 4-hydroxynonenal (4-HNE, a measure of oxidative damage), cytokeratin-7 (CK-7, a marker of precholangiocyte cellularity), or H2AX*γ* (a marker of DNA double strand breaks). Scoring of pathology was carried out using procedures described previously (Kleiner *et al.*, 2005; Lanaspa *et al.*, 2018; Monks *et al.*, 2018). 4-Hydroxynonenal IHC was performed using a 4-HNE antibody developed by the Petersen Laboratory (see Acknowledgements) (Shearn *et al.*, 2013). Cytokeratin-7 IHC was performed using a CK-7 antibody (ab181598, Abcam, Cambridge, MA). H2AX*γ* IHC was performed using a Phospho-Histone H2A.X antibody (Ser139) (Rabbit mAb #9718, Cell Signaling Technology, Danvers, MA, USA). Histologic images were captured on a microscope (Olympus BX51) equipped with a four-megapixel digital camera (Macrofire Optronics; Goleta, CA) using the PictureFrame Application 2.3 (Optronics). For quantification of the 4-HNE and CK-7 IHC, five 100x magnification field images were made in a tiling fashion across the stained section and the positive pixels in each image were quantified using “SlideBook” software (Intelligent Imaging Innovations, Denver, CO) (Petersen *et al.*, 2018). Quantification of H2AX*γ* IHC required counting positive cells in thirty consecutive 400x magnification fields. All IHC analyses were performed by an investigator blinded to the animal treatments. Hepatocytes were recognized by their size, shape, locale and nuclear size, shape and staining pattern. Non-hepatocytes were those cells in the liver tissue that were not hepatocytes; these included predominately endothelial cells, Kupffer cells, and stellate cells. Differences between control, 500 and 5,000 mg/L 1,4-DX-exposed mice were determined using one-way ANOVA with Tukey’s correction for multiple comparisons. p<0.05 was considered statistically significant. Students unpaired t-tests were used to compare results in mice exposed to 5,000 mg/L 1,4-DX for four weeks with those in control mice. Due to positive staining of H2AX*γ* in liver, H2AX*γ* staining was also carried out on formalin-fixed, paraffin-embedded kidney tissues from mice exposed to 0, 500, or 5,000 mg/L 1,4-DX for one week, and 0 or 5,000 mg/L 1,4-DX for four weeks, to determine the systemic effect of 1,4-DX on tissue DNA damage.

### 2.4 Hepatic transcriptome analyses

Liver tissues from mice exposed to 0 (control) or 5,000 mg/L 1,4-DX in drinking water for four weeks (n=3/group) were submitted to the Yale Center for Genome Analysis for RNA-sequencing (RNA-Seq) analysis.

#### RNA-Seq library preparation

mRNA (≈500 ng total) was purified with oligo-dT beads and sheared by incubation at 94°C (first-strand synthesis with random primers). Second strand synthesis was performed using dUTP to generate strand-specific sequencing libraries. The cDNA library was then end-repaired, A-tailed adapters were ligated, and second-strand digestion was performed using uracil-DNA-glycosylase. Indexed libraries (that met appropriate cut-offs for both) were quantified by qRT-PCR using a commercially-available kit (KAPA Biosystems), and insert size distribution was determined using a LabChip GX or Agilent Bioanalyzer. Samples with a yield ≥0.5 ng/μL were used for sequencing.

#### Flow cell preparation and sequencing

Sample concentrations were normalized to 10 nM and loaded onto Illumina Rapid or High-output flow cells at a concentration that yielded 150-250 million passing filter clusters per lane. Samples were sequenced using 75 bp single or paired-end sequencing on an Illumina HiSeq 2000 or 2500 according to manufacturer’s protocols. The 6 bp index was read during an additional sequencing read that automatically followed the completion of read 1. Data generated during sequencing runs were simultaneously transferred to the YCGA high-performance computing cluster. A positive control (prepared bacteriophage Phi X library) provided by Illumina was spiked into every lane (at a concentration of 0.3%) to monitor sequencing quality in real time.

#### Transcriptomic data analysis

Statistical analyses were performed using Partek Flow software (version 7.0, build 7.18.0130.2018 Partek Inc., St. Louis, MO, USA). Paired-end reads were trimmed using a base quality score threshold of >20 and aligned to the Genome Reference Consortium Mouse Build 38 (mm10) with the STAR 2.5.3a aligner. Quantification was carried out using the transcript model Ensembl Transcripts release 91. Differential expression analysis was performed using the R package DESeq2 which provides a quantitative analysis using shrinkage estimators for dispersion and fold-change (Love *et al.*, 2014). Fisher’s Exact test was used to calculate fold-change. Genes in 1,4-DX-treated mice with an absolute fold-change of 1.5 and Benjamini-Hochberg false discovery rate (FDR) of p<0.05 relative to control mice were considered as differentially-expressed genes (DEGs). A negative fold-change indicates down-regulation of the DEG in 1,4-DX-treated mice compared to controls. Qlucore Omics Explorer 3.4 (Qlucore AB, Lund, Sweden) was used for principal components analysis (PCA) and hierarchical clustering analysis (HCA) heatmap generation of log2-transformed global expression values. Raw and processed RNA-seq data was submitted to Gene Expression Omnibus (GEO) database: GSE158087.

#### Pathway analysis

Network analysis was performed using Ingenuity Pathway Analysis software (IPA, QIAGEN Redwood City, USA, www.qiagen.com/ingenuity) to identify pathways, diseases and functions over-represented in the DEGs. In this manually-curated knowledge base, each gene symbol was mapped to its corresponding gene object in the Ingenuity Pathway Knowledge Base. Significant (Benjamini-Hochberg FDR p<0.05) pathway enrichment within a reference network was performed using Fisher’s Exact test.

### 2.5 Metabolomic analyses

#### 2.5.1 Sample preparation

##### Liver and kidney tissue samples

Samples (50.00 ± 2.50 mg each) were homogenized with ceramic beads in 100 μL H2O for three cycles (20 sec at 6,000 rpm, with 5 sec between each cycle) using a Precellys Evolution homogenizer (Bertin Technologies SAS, Montigny-Le-Bretonneux, France). Dry ice was used to keep the samples at a low temperature during homogenization using a Cryolys attachment (Bertin Technologies SAS, Montigny-Le-Bretonneux, France). The tissue homogenates (100 μL aliquots) were snap frozen using liquid nitrogen and stored at −80°C for later use. For metabolomic analyses, samples were thawed on ice and 400 μL of ice-cold methanol:acetonitrile (1:1, v/v) was added. The resulting solution was vortexed for 30 sec, and centrifuged at 15,000 rpm for 15 min at 4°C. The resulting supernatant was placed in a clean tube, incubated at −20°C for 1 h (to further precipitate proteins), and subsequently subjected to centrifugation at 15,000 rpm for 15 min at 4°C. The supernatant was evaporated to dryness in a vacuum concentrator (ThermoFisher Scientific, Waltham, MA). Dry extracts were reconstituted in 150 μL acetonitrile:water (1:1, v/v), and centrifuged at 15,000 rpm for 10 min at 4°C (to remove insoluble debris). The resulting supernatant was transferred to liquid chromatography-mass spectrometry (LC-MS) glass vials for LC-MS analysis. A quality control (QC) sample was prepared by pooling a small aliquot from each reconstituted sample.

##### Urine samples

*S*amples were thawed overnight in a refrigerator and prepared using solid phase extraction (SPE). A 96 well Oasis HLB μElution Plate (Waters Corporation, Milford, MA) was conditioned with 200 μL methanol and then 200 μL of each sample (urine:water 1:4 v/v) was loaded onto the plate by pipette. The samples were extracted using an Oasis 96-well Extraction Plate Manifold connected to a vacuum pump, and the eluate subsequently transferred to a 96-well collection plate (Waters Corporation, Milford, MA) (containing samples for hydrophilic interaction LC-MS (HILIC-MS)). Methanol (200 μL) was then added twice onto the original μElution Plate using a multichannel pipette and the eluate collected onto an additional 96-well collection plate (containing samples for reversed-phased LC-MS (RPLC-MS)). All collection plate samples were then concentrated to dryness by speed vacuum (SPD 111, Thermo Scientific, Waltham, MA) at room temperature, reconstituted in 150 μL acetonitrile: water (1:1, v/v), and centrifuged at 15,000 rpm for 10 min at 4°C (to remove insoluble debris). The resulting supernatant was transferred to liquid chromatography-mass spectrometry (LC-MS) glass vials for LC-MS analysis. A QC sample was prepared by pooling a small aliquot from each reconstituted sample.

##### Fecal samples

Frozen fecal pellets (25 mg) were homogenized in 200 μL H2O for three cycles (20 sec at 5,500 rpm, with 5 sec between each cycle) using a Precellys Evolution homogenizer (Bertin Technologies SAS, Montigny-Le-Bretonneux, France). Dry ice was used to keep the samples at a low temperature during homogenization using a Cryolys attachment (Bertin Technologies SAS, Montigny-Le-Bretonneux, France). Samples (150 μL) were further extracted in 150 μL methanol/acetonitrile (1:1), vortexed for 30 sec, and subjected to sonication in an ultrasonic waterbath (FisherBrand™, Thermo Scientific, Waltham, MA) for 10 min at 4°C. Samples were then incubated for 2 h at −20°C, and centrifuged at 13,000 rpm for 15 min at 4°C. The resulting supernatant was transferred into a 1.5 mL tube and evaporated to dryness in a vacuum concentrator (SPD 111, Thermo Scientific, Waltham, MA). Dry extracts were reconstituted in 100μL acetonitrile: water (1:1, v/v), vortexed for 30 sec, and sonicated for 10 min at 4°C. Samples were then centrifuged at 13,000 rpm for 15 min and 4°C. The resulting supernatant fluid was obtained and diluted with 400 μL acetonitrile: water (1:1, v/v), for subsequent LC-MS analysis. A QC sample was prepared by pooling a small aliquot from each reconstituted sample.

#### 2.5.2 Untargeted metabolomic analysis of liver, kidney, urine and feces samples

All extracted samples were analyzed on a quadrupole time-of flight (Q-ToF) mass spectrometer (Xevo G2-XS Q-ToF, Waters Corporation, Milford, MA) equipped with an ultra-performance liquid chromatography (UPLC) Acquity I Class (Waters Corporation, Milford, MA) unit. Chromatographic separation was performed using an Acquity UPLC BEH C18 column (particle size, 1.7 μm; 50 mm (length) × 2.1 mm (i.d.)) (Waters Corporation) equipped with a BEH C18 VanGuard pre-column (particle size, 1.7 μm; 50 mm (length) × 2.1 mm (i.d.)) (Waters Corporation, Milford, MA) for RPLC-MS. An Acquity BEH Amide column (particle size, 1.7 μm; 100 mm (length) × 2.1 mm (i.d.)) (Waters Corporation, Milford, MA) equipped with a BEH Amide VanGuard pre-column (5 x 2.1 mm, i.d.; 1.7μm) (Waters Corporation, Milford, MA) was used for chromatographic separation for HILIC-MS. The mobile phase for RPLC-MS analysis consisted of A (0.1% formic acid in water) and B (0.1% formic acid in acetonitrile) delivered at a flow rate of 0.5 mL/min. The linear gradient elution started at 1% B (0-1 min), 1%-100% B (1-8 min), 100% B (8-10 min), 100-1% B (10.0-10.1 min) and continuing at 1% B (10.1-12.0 min). The mobile phase for HILIC-MS analysis consisted of A (25 mM ammonium hydroxide and 25 mM ammonium acetate in water) and B (acetonitrile) delivered at a flow rate of 0.5 mL/min. The linear gradient elution started at 95% B (0-0.5 min), 95%-65% B (0.5-7 min), 65-40% B (7-8 min), 40% B (8-9 min), 40-95% B (9-9.1 min) and continuing at 95% B (9.1-12.0 min). The injection volume for all samples and standard solutions was 1 μL. QC samples were analyzed every five to eight injections. The column temperature was set at 25°C for RPLC and 30°C for HILIC, and the sample tray temperature was maintained at 4°C. For MS analyses, the electrospray ionization source (ESI) was operated in positive and negative mode. Q-ToF-MS scan data (300 ms/scan; mass scan range 50-1200 Da) were first acquired for each biological sample. Thereafter, MS^e^ fragmentation data were acquired for metabolite identification (low energy scan: 200 ms/scan, collision energy 6 eV; high energy scan: 100 ms/scan, collision energy 40 eV, mass scan range 25-1000 Da). ESI source parameters were as follows: 1.8 kV capillary voltage, 40 V sampling cone, 50°C source temperature, 550°C desolvation temperature, 40 L/hr cone gas flow, 900 L/hr desolvation gas flow. Of note, for each sample type (liver, kidney, urine or stool), a QC sample for each sample type was injected 15 times before the individual samples for that sample type were injected, to ensure matrix stabilization.

##### Data analysis

Raw MS data (.raw) files were converted to mzML format (ProteoWizard version 3.06150), and R-XCMS (version 3.0.2) was used for data processing. The generated data matrix consisted of the mass-to-charge ratio (*m*/*z*) value, retention time (RT), and peak abundance. The workflow involved typical data-processing steps, including peak detection and peak alignment.

Final peak picking parameters were: peak width = c(5,30); snthresh = 5; mzdiff = 0.05; ppm = 15; alignment (bw = 10, mzwid = 0.05); and retention time correction (obiwarp, plottype = c(deviation), profstep = 1). The R package “CAMERA” was used for peak annotation after XCMS data processing. The minfrac parameter in XCMS was set to 0.5 to discard any metabolites that did not appear in at least 50 % of the quality control (QC) samples; this was performed to eliminate analytical noise. Subsequently, the R package “MetNormalizer” was applied using support vector regression (SVR) normalization to remove the intra batch variation (Shen *et al.*, 2016; Shen *et al.*, 2019). Missing values were imputed using the k-nearest neighbor method (k=5). The data resulting from these analyses were subjected to multivariate (prcomp; i.e., principal components analysis) and univariate (wilcox.test, p.adjust) analyses using the R platform (version 3.4.3). The Wilcoxon rank sum test with Benjamini-Hochberg FDR correction (*α* = 5%) was used to detect significant differences between samples from the 1,4-DX-treated and control mice. To construct the volcano plots, fold-change and p values were log2- and −log10-transformed using R function “log ()”. Function “plot()” was modified to plot the log2-transformed fold-change against −log10-transformed p value for each variable. Metabolomics data are available at the NIH Metabolomics workbench (project number PR001030, http://www.metabolomicsworkbench.org/).

##### Putative annotation

For metabolite annotation, the Metabolite annotation and Dysregulated Network Analysis (MetDNA) workflow was used. This uses a metabolic reaction network (MRN)-based recursive algorithm for metabolite annotation (Shen and Zhu, 2019).

#### 2.5.3 Targeted analysis of liver and fecal bile acids

As our transcriptomic data and pathway analysis revealed possible perturbations in bile acid metabolism, bile acids were targeted for analysis by Q-ToF-MS multiple reaction monitoring (MRM). This type of analysis can quantify bile acids with target enhancement, i.e., a precursor ion is selected by the quadrupole and fragmented in the collision cell. The Q-ToF pusher was synchronized with the mass-to-charge ratio (m/z) of the precursor or a product ion; this maximized the duty cycle for a target m/z range and increased the response and selectivity.

##### LC-MS analysis

The liver and stool samples from control mice and mice exposed to 5,000 mg/L 1,4-DX for one and four weeks were used for bile acid quantification. Samples that had been previously extracted for untargeted metabolomics were further analyzed using MRM on the same Q-ToF mass spectrometer (Xevo G2-XS Q-ToF, Waters Corporation, Milford, MA). Chromatographic separation was performed using an Acquity UPLC BEH C18 column (particle size, 1.7 μm; 50 mm (length) × 2.1 mm (i.d.)) (Waters Corporation, Milford, MA) equipped with a BEH C18 VanGuard pre-column (particle size 1.7μm; 50 mm (length) x 2.1 mm (i.d.)) (Waters Corporation, Milford, MA). The mobile phase consisted of A (aqueous buffer containing 1 mM ammonium formate and formic acid (pH 4.39)) and B (acetonitrile/isopropanol (1:1 v/v)) delivered at a flow rate of 0.4 mL/min. The linear gradient elution started at a ramp of 20-30% B (0-3 min), 30-40% B (3-4 min), 50-70% B (4-5 min), 70-90% B (5.0-5.2 min), continuing at 90% B (5.2-6 min), ramp of 90-20% B (6.0-6.1 min) and continuing at 20% B (6.1-8.0 min). The injection volume for all samples and standard solutions was 2 μL. The column temperature was set at 55°C, and the sample tray temperature was maintained at 8°C. For MS analysis, an electrospray ionization source was operated in negative mode (ESI−) and MassLynx 4.1.0 software (Waters Corporation, Milford, MA) was used to acquire the data. The ESI source parameters were as follows: 2 kV spray voltage, 30 V cone voltage, 120°C source temperature, 500°C desolvation temperature, 50 L/h cone gas flow, 900 L/h desolvation gas flow. The Q-ToF MRM was adopted for enhanced target quantitative analysis according to the transitions and collision energy levels described for each analyte (as listed in Table 1).

**Table 1:** Multiple reaction monitoring (MRM) transitions, retention times and collision energy levels for the 19 bile acids that were quantified in the liver samples.

| Bile acid | Precursor ion<sup>A</sup><br>(M-H)<sup>-</sup> | Product ion<sup>B</sup> | Retention time<sup>C</sup><br>(min) | Collision energy<sup>D</sup><br>(eV) |
| --- | --- | --- | --- | --- |
| 7-ketodeoxycholate (7-keto-DCA ) | 405.2646 | 405.2646 | 3.81 | 0 |
| 7-ketolithocholic acid (7-keto-LCA ) | 389.2697 | 389.2697 | 4.47 | 0 |
| Chenodeoxycholic acid (CDCA) | 391.2853 | 391.2853 | 4.85 | 0 |
| Cholic acid (CA) | 407.2802 | 407.2802 | 4.4 | 0 |
| Deoxycholic acid (DCA) | 391.2853 | 391.2853 | 4.93 | 0 |
| Glycochenodeoxycholic acid (GCDCA) | 448.3068 | 74.0244 | 4.22 | -40 |
| Glycocholic acid (GCA) | 464.3017 | 74.0244 | 3.55 | -55 |
| Glycodeoxycholic acid (GDCA) | 448.3068 | 74.0244 | 4.33 | -40 |
| Glycohyodeoxycholic acid (GHDCA) | 448.3068 | 74.0244 | 3.43 | -40 |
| Glycolithocholic acid (GLCA) | 432.3119 | 74.0244 | 4.69 | -40 |
| Glycoursodeoxycholic acid (GUDCA) | 448.3068 | 74.0244 | 3.22 | -40 |
| Hyodeoxycholic acid (HDCA) | 391.2853 | 391.2853 | 4.46 | 0 |
| Lithocholic acid (LCA) | 375.2904 | 375.2904 | 5.31 | 0 |
| Taurochenodeoxycholic acid (TCDCA) | 498.2894 | 79.9568 | 4.11 | -70 |
| Taurocholic acid (TCA) | 514.2843 | 79.9568 | 3.30 | -65 |
| Taurodeoxycholic acid (TDCA) | 498.2894 | 79.9568 | 4.23 | -70 |
| Taurohyodeoxycholic acid (THDCA) | 498.2894 | 79.9568 | 3.30 | -65 |
| Taurolithocholate (TLCA) | 482.2946 | 79.9568 | 4.57 | -70 |
| Ursodeoxycholic acid (UDCA) | 391.2853 | 391.2853 | 4.38 | 0 |

<sup>A</sup> Precursor ion; m/z of bile acid before fragmentation
<sup>B</sup> Product ion; ion resulting from fragmentation of precursor ion, that enables identification of metabolite
<sup>C</sup> Retention time; time in which the bile acid elutes from the column
<sup>D</sup> Collision energy; energy provided to fragment the precursor ion.

##### Standard solutions preparation

Standard stock solutions (1 mg/mL) of bile acids were individually prepared by dissolving each bile acid in methanol:water (1:1 v/v). All 19 bile acids were combined in a pooled solution at 5 μg/mL in acetonitrile:water (1:1 v/v). This solution was then serially diluted with acetonitrile:water (1:1) to obtain standard working solutions containing 1.6, 5.0, 16, 50, 160 and 500 ng/mL of cholic acid (CA), deoxycholic acid (DCA), hyodeoxycholic acid (HDCA), lithocholic acid (LCA), and taurolithocholate (TLCA). The working solutions 1.6, 5.0, 16, 50, 160, 500, and 1600 ng/mL of 7-ketodeoxycholate (7-keto-DCA), 7-ketolithocholic acid (7-keto-LCA), chenodeoxycholic acid (CDCA), glycochenodeoxycholic acid (GCDCA), glycocholic acid (GCA), glycodeoxycholic acid (GDCA), glycohyodeoxycholic acid (GHDCA), glycolithocholic acid (GLCA), glycoursodeoxycholic acid (GUDCA), taurochenodeoxycholic acid (TCDCA), taurocholic acid (TCA), taurodeoxycholic acid (TDCA), taurohyodeoxycholic acid (THDCA), and ursodeoxycholic acid (UDCA).

##### Data acquisition and processing

Data were acquired using MassLynx version 4.1 (Waters Corp., Milford, MA). TargetLynx software (Waters Corp., Milford, MA) was used to quantify bile acids by integrating peak areas of extracted ion chromatograms. The concentration of each metabolite was calculated by comparison to a standard curve of an authentic standard. Differences between control mice and 1,4-DX-exposed mice for individual bile acids were determined using Student’s unpaired t-test with Benjamini, Krieger and Yekutieli correction for multiple comparisons (Benjamini and Hochberg, 1995) (GraphPad Prism version 8.0 for Windows, GraphPad Software, La Jolla California USA). p < 0.05 was considered significant.

## Results

### 3.1 Effect of 1,4-DX exposure on body weight, water consumption, and average exposure

All mice receiving doses of 1,4-DX (0, 50, 500 or 5,000 mg/L) for one or four weeks were healthy and active during the course of the experiments. None of the mice (control or 1,4-DX-exposed) showed any overt adverse health effects. In addition, control and 1,4-DX-exposed mice did not exhibit any differences (p>0.05) in body weight, liver weights or daily water consumption (**Supplementary Figure S1**). The terminal body weights, liver weights or relative liver weight (%), daily water intake, and data on the 1,4-DX intake (daily dose or total dose (mg/kg/bw)) for one week or four weeks are presented in **Supplementary Table 1**. The average daily and total intake of 1,4-DX increased in proportion to the drinking water concentration and the duration of 1,4-DX exposure (**Supplementary Table 1**).

### 3.2 Histopathology

Immunohistochemical analysis of 4-HNE expression in the liver from 1,4-DX-treated mice revealed only a low-level signal, and no differences were observed between the control and 1,4-DX-exposed mice (500, 5000 mg/L for one or four weeks; data not shown).

H2AX*γ* expression in liver was low, i.e., <1 positive hepatocyte per 400x field (**Figure 1A**). As a result, thirty fields per sample were counted, and positive cells were recorded according to expression in hepatocytes or non-hepatocytes (i.e., Kupffer cells, lymphocytes, endothelial cells, stellate cells and cholangiocytes). Hepatocellular carcinoma is derived from hepatocytes or their precursors; therefore, hepatocytes and non-hepatocytes were examined separately. After one week of exposure, H2AX*γ* expression in hepatocytes was not different between control mice and those exposed to 500 mg/L 1,4-DX. By contrast, exposure to 5,000 mg/L 1,4-DX for one week doubled the number of hepatocytes expressing H2AX*γ* (**Figure 1A, B**); this increase also occurred in the livers of mice exposed to 5,000 mg/L 1.4-DX for four weeks (livers from mice exposed to 500 mg/L 1,4-DX for four weeks were not stained for H2AX*γ*) (**Figure 1B**). For non-hepatocytes, H2AX*γ* was significantly increased in mice exposed for one week to 5,000 mg/L 1,4-DX but not to 500 mg/L 1,4-DX. After four weeks exposure to 5,000 mg/L, H2AX*γ* was not increased (**Figure 1C**). The histology of the kidneys from the 1,4-DX-exposed mice revealed no significant differences in the number of H2AX*γ*-stained cells between control mice and any of the mice exposed to 1,4-DX (at any concentration or exposure duration) (data not shown).

**Figure 1:**
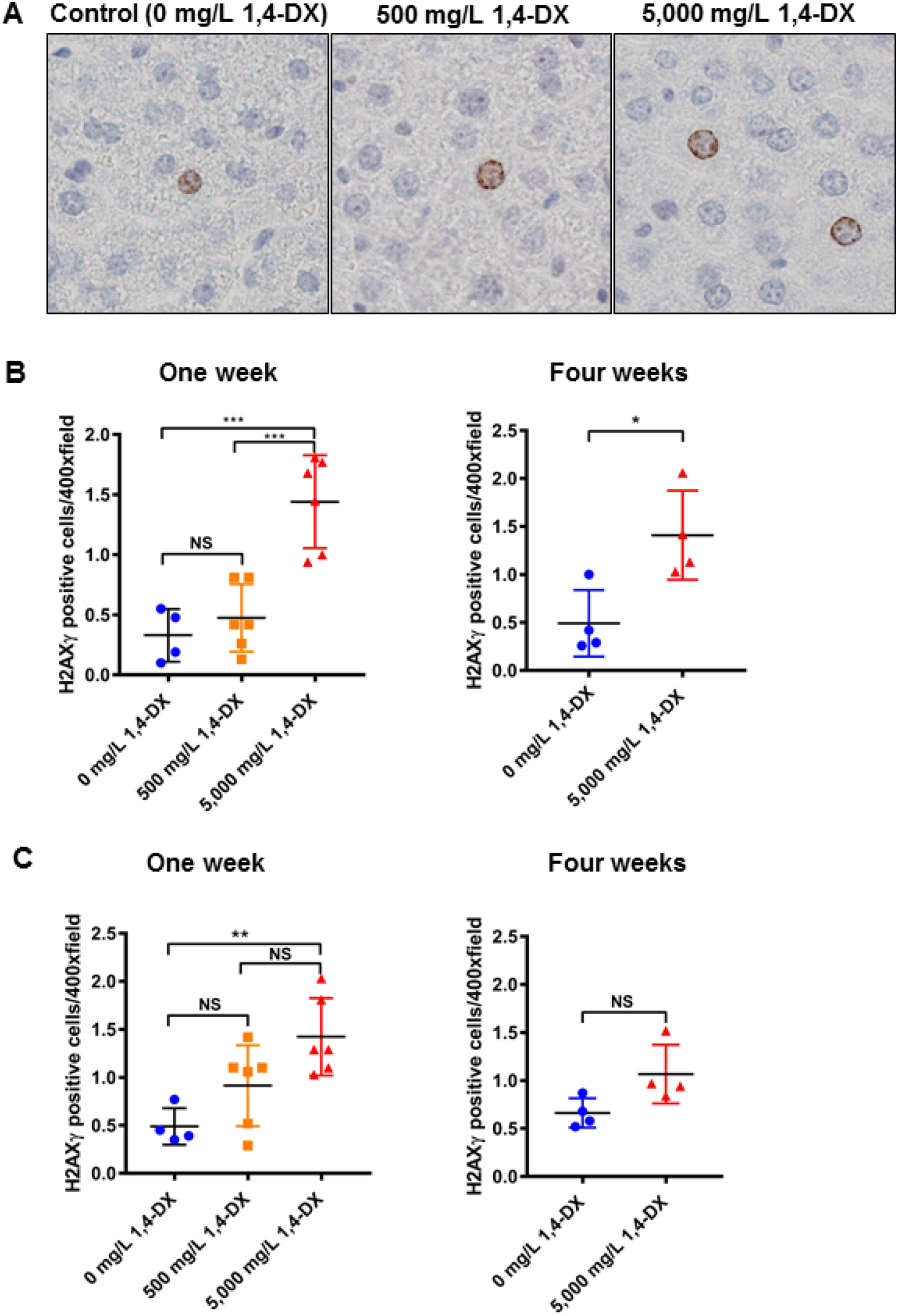
H2AXγ expression in the liver of mice exposed to 1,4-DX. **A)** Liver H2AXg hepatocyte staining after one week 1,4-DX exposure. Formalin-fixed, paraffin-embedded liver tissue was sectioned and immunostained for the presence of H2AXγ in control (untreated) mice and mice exposed to 1,4-DX (500 or 5,000mg/L) in drinking water for one week. Representative images (400x) are shown for hepatocyte cell staining in control (0 mg/L) 1,4-DX (left panel), 500 mg/L 1,4-DX-(middle panel) and 5,000 mg/L 1,4-DX-(right panel) exposed mice. **B)** Liver H2AXg hepatocyte quantification. The number of hepatocytes staining positively for H2AXγ in thirty consecutive 400x magnification fields per liver. **C)** Liver H2AXg non-hepatocyte quantification. The number of non-hepatocyte cells staining positively for H2AXγ in thirty consecutive 400x magnification fields per liver. Data are presented as individual values and the means ± SD for each treatment group (n=4-6 mice/group). For one-week exposure, differences between groups of mice were determined using one-way ANOVA with Tukey correction for multiple comparisons. For four weeks exposure, differences between groups of mice were determined using a Student’s unpaired t-test. NS = not significant; *p<0.05, ** p<0.01, *** p<0.001

The liver can sense damage and immediately start to repopulate itself. Therefore, CK-7 immunostaining was used to identify any changes in the population of precursor cells, i.e., precholangiocytes. After one week of exposure to 1,4-DX, there was no difference between CK-7-positive cells in control and 1,4-DX-exposed (500 or 5,000 mg/L) mice. After four weeks of 5,000 mg/L 1,4-DX exposure, the population of CK-7-positive cells increased by 20%, and such cells were present in the correct histological locale for precholangiocytes, i.e., periductularly and spreading out around the portal vein (**Figure 2A, B**). Of note, the CK-7 immunostaining can also identify cholangiocytes but there was no increase in these cells. CK-7 staining was not carried out in kidney because the staining does not label the precursor cell population in this tissue.

**Figure 2:**
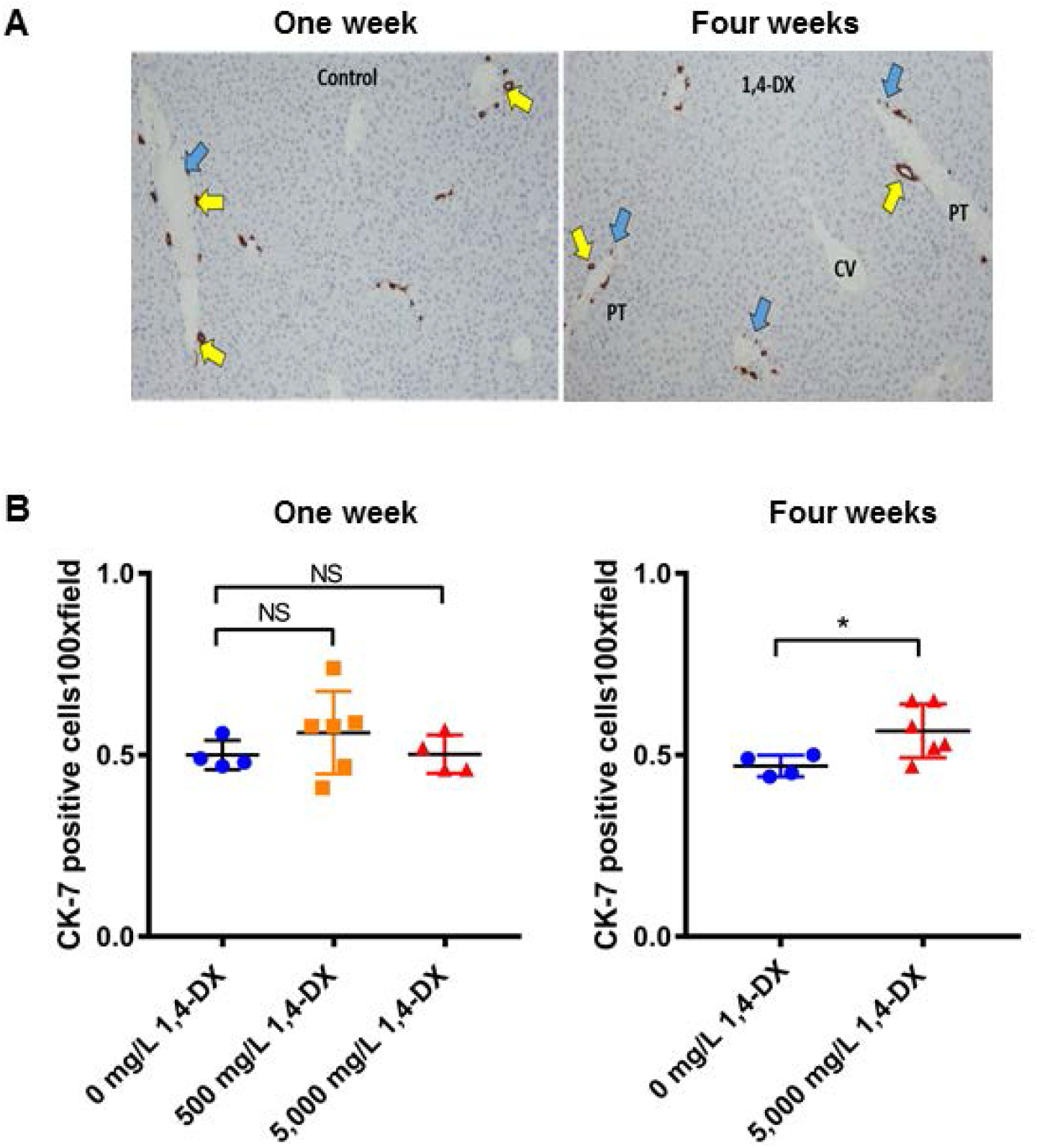
Cytokeratin-7 (CK-7) expression in the liver of mice exposed to 1,4-DX. **A)** Representative images of tissue sections from liver tissue obtained from mice exposed to 0 (control), 500 or 5,000 mg/L 1,4-DX for one week (left panel) or to 0 (control) or 5,000mg/L 1,4-DX for four weeks (right panel). Precholangiocytes associated with bile ducts (yellow arrows) and single precholangiocytes close to the portal triad (PT) (blue arrows) are highlighted. Magnification: 100x. **B)** The number of cells staining positively for CK-7 in five consecutive 100x magnification fields per liver. Data are presented as individual values and the means ± SD for each treatment group (n=4-6 mice/group). For one-week exposure, differences between groups of mice were determined using a one-way ANOVA with Tukey correction for multiple comparisons. For four weeks exposure, differences between groups of mice were determined using a Student’s unpaired t-test. NS = not significant; *p<0.05.

### 3.3 Hepatic transcriptome analysis

RNA-Seq was performed on liver tissue samples from three mice exposed to 0 (control) or 5,000 mg/L 1,4-DX for four weeks. Principal components analysis (PCA) showed separation of control and 1,4-DX-exposed mice, indicating differences in the hepatic transcriptome (**Figure 3A)**. Further analysis by hierarchical clustering analysis identified 65 differentially-expressed transcripts in the livers of mice exposed to 1,4-DX with a FDR-corrected p-value < 0.05 and fold-change >1.5 (**Figure 3B, Table 2)**. *Cyp3a16* (human orthologue *CYP3A5*) was highly down-regulated (fold-change=95.6) (**Table 2**). Using these differentially-expressed genes, pathway analysis was implemented to identify the molecular pathways enriched within the livers of 1,4-DX-exposed mice. Of the differentially-expressed genes, 58 were mapped to functional pathways. The genes from the mice were converted into the human orthologues to enable further understanding of the potential impact of 1,4-DX exposure in humans.

**Table 2.**
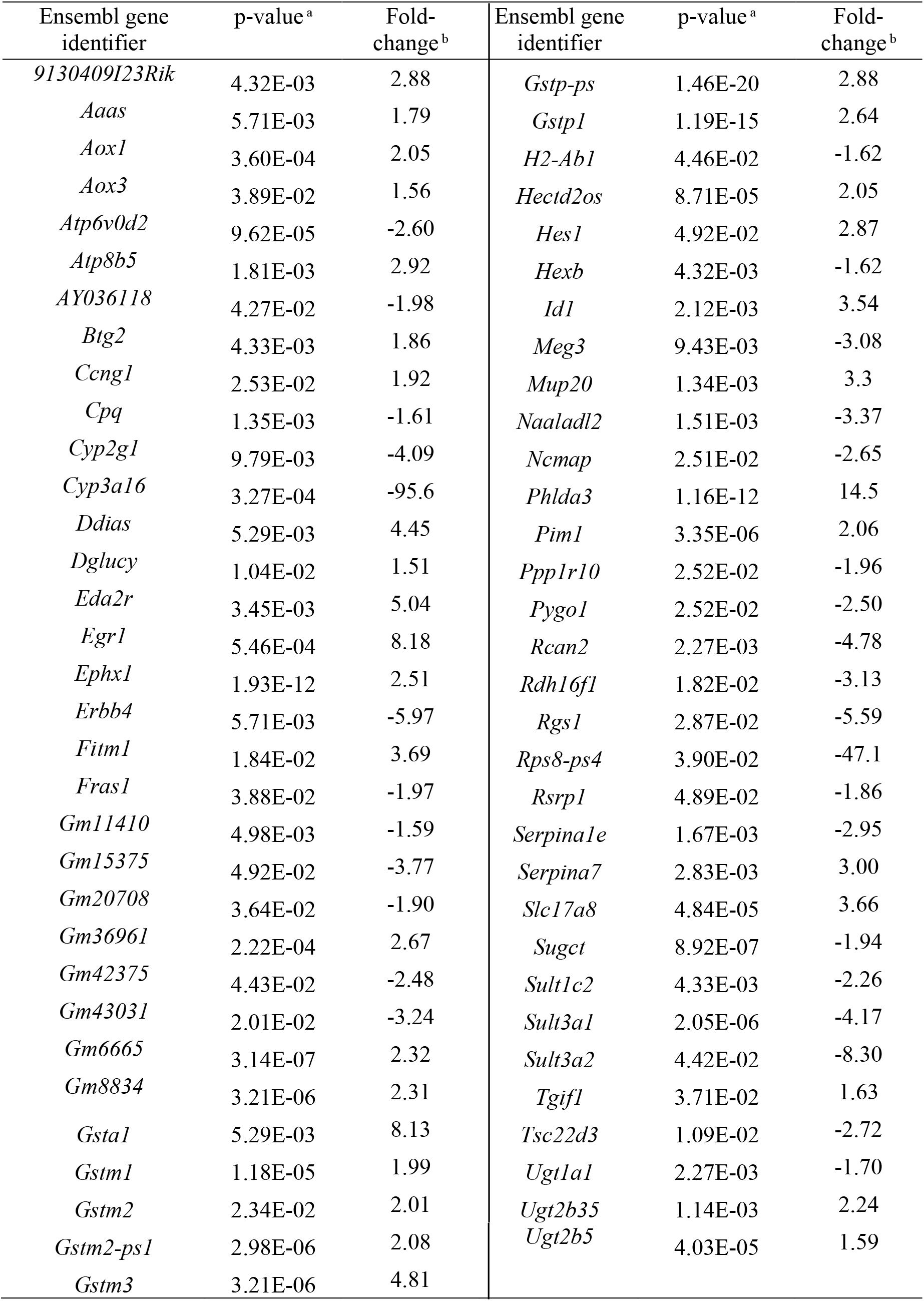
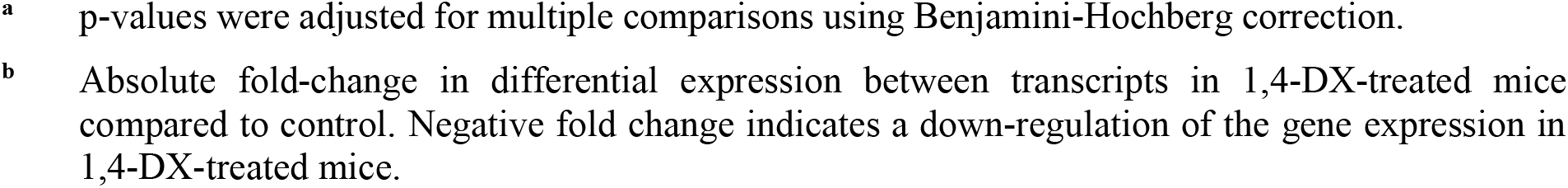
Differentially expressed genes in the liver of mice exposed to 5,000 mg/L 1,4-DX in drinking water for 4 weeks compared to control mice (genes shown have a p-value < 0.05, fold-change > 1.5 between control and 1,4-DX-exposed mice).

**Figure 3:**
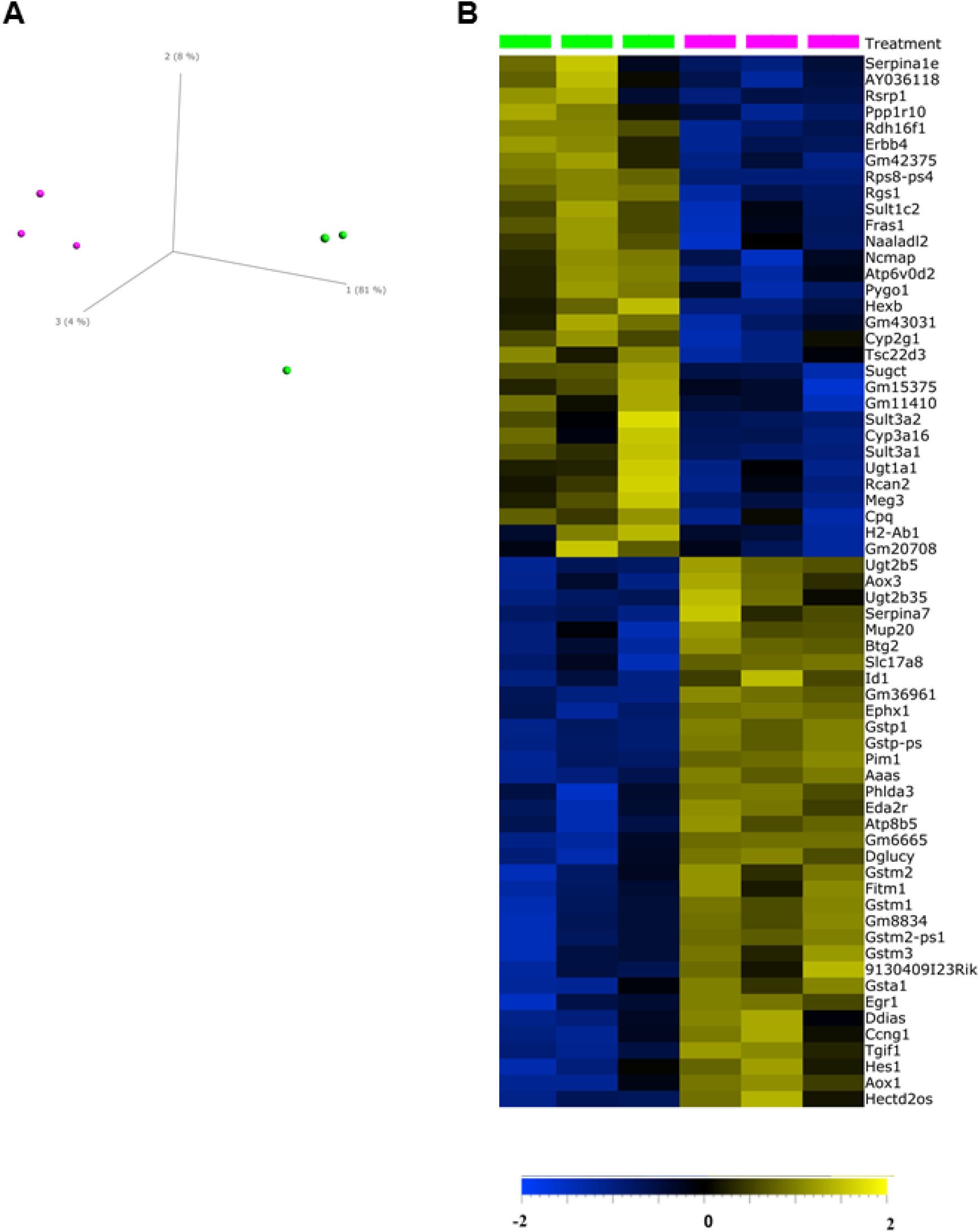
Transcriptomics changes in the liver of 1,4-DX-exposed mice. **A)** Principal components analysis scores plot of transcriptomics data from liver samples of mice exposed to 0 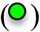 or 5,000 mg/L 1,4-DX in water 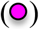 for four weeks. **B)** Hierarchical clustering analysi*s* heatmap showing the 65 differentially-expressed (FDR-corrected p-value > 0.05 and log2 fold-change >1.5) genes in livers of mice exposed to 0 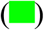 and or 5,000 mg/L 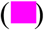 1,4-DX for four weeks. The log2 fold-change in expression in 1,4-DX-exposed mice are shown on a scale from −2.0 (blue) to 2.0 (gold)

The enriched metabolic pathways were (in order of descending levels of significance): xenobiotic metabolism signaling, nicotine degradation III, glutathione-mediated detoxification, nicotine degradation II, LPS/IL-1 mediated inhibition of RXR function, NRF2-mediated oxidative stress response, aryl hydrocarbon receptor signaling, melatonin degradation I, superpathway of melatonin degradation, thyroid hormone metabolism II, aryl hydrocarbon receptor signaling, serotonin degradation, PXR/RXR activation, glutathione redox reactions I, dermatan sulfate biosynthesis (late stages), chondroitin sulfate biosynthesis (late stages) and dermatan sulfate biosynthesis (**Table 3**). The PXR/RXR pathway had two out of the four identified genes down-regulated, while LPS/IL-1-mediated inhibition of RXR function showed *CYP3A5, SULT2C2, SULT3A1*, and *SULT3A2* to be down-regulated. Glutathione S-transferase genes were up-regulated (**Table 3**)

**Table 3:** Canonical pathways enriched in livers from mice exposed to 5,000 mg/L 1,4-DX in drinking water for 4 weeks. Differentially-expressed genes were subjected to pathway analysis using IPA. Gene names are shown as human orthologs.

| Canonical pathways | Adjusted p-value (FDR)<sup>a</sup> | Genes enriched within pathway |
| --- | --- | --- |
| Xenobiotic Metabolism Signaling | 7.08E-09 | *CYP3A5, GSTA5, GSTM1, GSTM3, GSTM5, GSTP1, SULT1C2, SULT3A1/SULT3A2, UGT1A1, UGT2B28, UGT2B7* |
| Nicotine Degradation III | 1.78E-07 | *AOX1, AOX3, CYP3A5, UGT1A1, UGT2B28, UGT2B7* |
| Glutathione-mediated Detoxification | 2.69E-07 | *GSTA5, GSTM1, GSTM3, GSTM5, GSTP1* |
| Nicotine Degradation II | 2.69E-07 | *AOX1, AOX3, CYP3A5, UGT1A1, UGT2B28, UGT2B7* |
| LPS/IL-1 Mediated Inhibition of RXR Function | 8.91E-07 | *CYP3A5, GSTA5, GSTM1, GSTM3, GSTM5, GSTP1, SULT1C2, SULT3A1/SULT3A2* |
| NRF2-mediated Oxidative Stress Response | 6.03E-06 | *AOX1, EPHX1, GSTA5, GSTM1, GSTM3, GSTM5, GSTP1* |
| Melatonin Degradation I | 6.03E-06 | *CYP3A5, SULT1C2, UGT1A1, UGT2B28, UGT2B7* |
| Superpathway of Melatonin Degradation | 7.59E-06 | *CYP3A5, SULT1C2, UGT1A1, UGT2B28, UGT2B7* |
| Thyroid Hormone Metabolism II (via Conjugation and/or Degradation) | 3.39E-05 | *SULT1C2, UGT1A1, UGT2B28, UGT2B7* |
| Aryl Hydrocarbon Receptor Signaling | 1.99E-04 | *GSTA5, GSTM1, GSTM3, GSTM5, GSTP1* |
| Serotonin Degradation | 2.82E-04 | *SULT1C2, UGT1A1, UGT2B28, UGT2B7* |
| PXR/RXR Activation | 3.89E-03 | *CYP3A5, GSTM1, UGT1A1* |
| Glutathione Redox Reactions I | 1.09E-02 | *GSTM1, GSTP1* |
| Dermatan Sulfate Biosynthesis (Late Stages) | 3.63E-02 | *SULT1C2, SULT3A1/SULT3A2* |
| Chondroitin Sulfate Biosynthesis (Late Stages) | 3.89E-02 | *SULT1C2, SULT3A1/SULT3A2* |
| Chondroitin Sulfate Biosynthesis | 4.90E-02 | *SULT1C2, SULT3A1/SULT3A2* |
| Dermatan Sulfate Biosynthesis | 4.90E-02 | *SULT1C2, SULT3A1/SULT3A2* |

<sup>a</sup> Fisher's Exact test p-value (adjusted for FDR) of differentially-expressed genes

We then examined links between the identified differentially-expressed genes in 1,4-DX-exposed mice and specific diseases and functions. Five differentially-expressed genes were linked to activated DNA damage response of cells and inhibited repair of DNA (**Figure 4**). Pleckstrin homology-like domain family A member 3 (*PHLDA3*), DNA damage-induced apoptosis suppressor (*DDIAS*), and glutathione S-transferase pi 1 (*GSTP1*), BTG anti-proliferation factor (*BTG2*) were up-regulated while protein phosphatase 1 regulatory subunit 10 (*PPPLR10*) was down-regulated (**Table 4**). Four genes were linked to hepatocellular carcinoma (**Figure 4**). In this pathway, the inhibitor of DNA binding 1 (*ID1*), aldehyde oxidase 1 (*AOX1*) and cyclin G1 (*CCNG1*) were up-regulated, and UDP glucuronosyltransferase family 1 member A1 (*UGT1A1*) was down-regulated. (**Table 4**).

**Table 4:** Categorization of differentially-expressed genes into diseases and functions. Differentially-expressed genes in mice exposed to 5,000 mg 1,4-DX for four weeks (from Table 2) were subjected to Ingenuity pathway analysis.

| Disease/Functions | Fold-change (Log<sub>2</sub>)<sup>a</sup> | p-value<sup>b</sup> |
| --- | --- | --- |
| ***DNA damage response (p=8.1E-07) and DNA repair functions (p=1.00E-02)*** |  |  |
| pleckstrin homology like domain family A member 3 (*PHLDA3*) | 14.5 | 1.17E-12 |
| DNA damage induced apoptosis suppressor (*DDIAS*) | 4.45 | 5.30E-03 |
| glutathione S-transferase pi 1 (*GSTP1*) | 2.64 | 1.19E-15 |
| protein phosphatase 1 regulatory subunit 10 (*PPPLR10*) | -1.96 | 2.50E-02 |
| BTG anti-proliferation factor (*BTG2*) | 1.86 | 4.33E-033 |
| ***Hepatocellular carcinoma (p=2.20E-04)*** |  |  |
| inhibitor of DNA binding 1 (*ID1*) | 3.54 | 2.12E-03 |
| aldehyde oxidase 1 (*AOX1*) | 2.05 | 3.60E-04 |
| cyclin G1 (*CCNG1*) | 1.92 | 2.53E-02 |
| UDP glucuronosyltransferase family 1 member A1 (*UGT1A1*) | -1.70 | 2.27E-03 |

<sup>a</sup> Log<sub>2</sub>[(gene expression in the liver of 5,000 mg/L 1,4-DX-exposed mice)/(gene expression in the liver of 0 mg/L 1,4-DX-exposed mice)]. Negative fold change indicates a down-regulation of the gene expression in 1,4-DX-treated mice.
<sup>b</sup> Fisher's Exact test p-value (adjusted for FDR) of differentially-expressed genes

**Figure 4:**
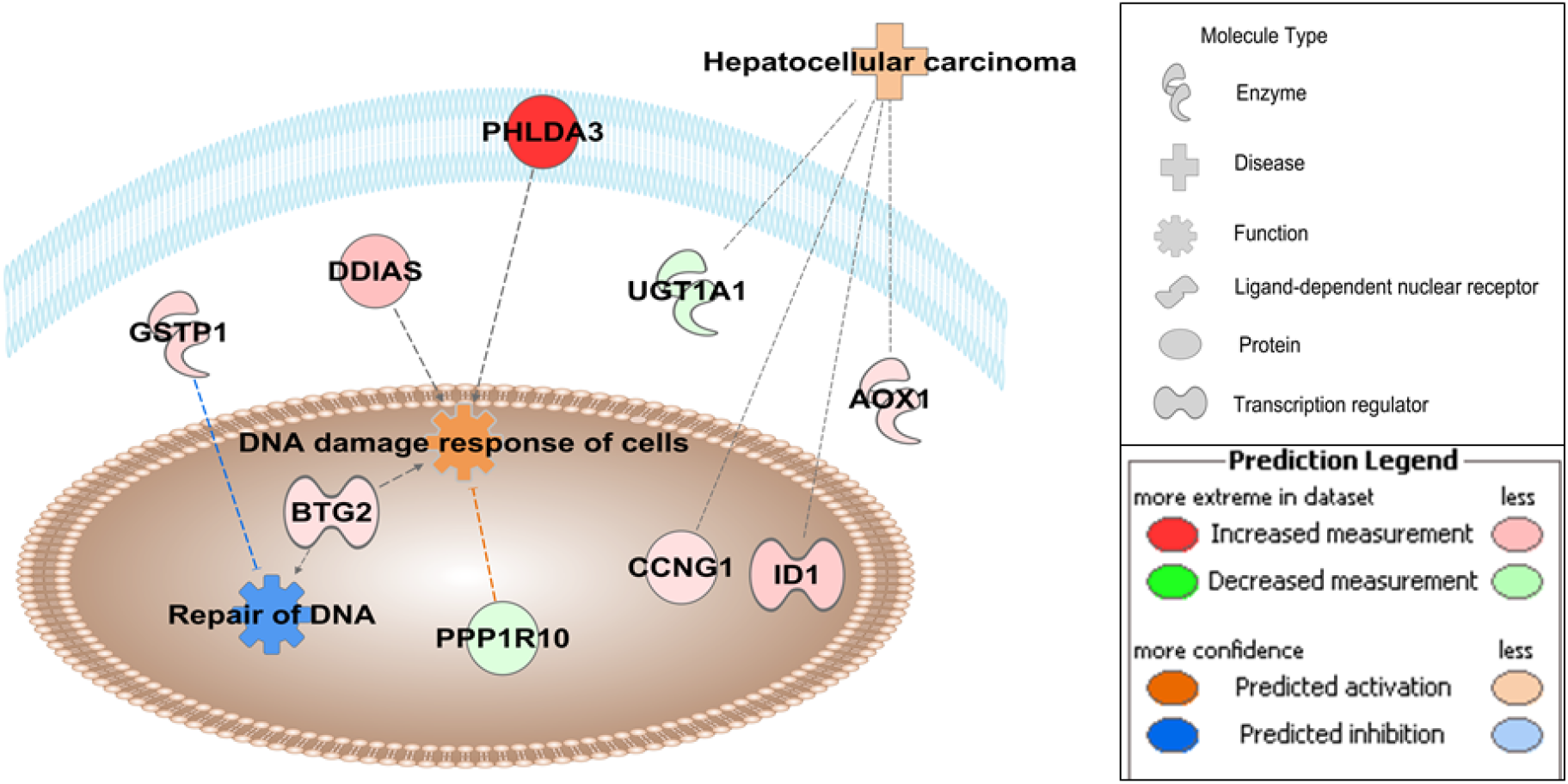
Pathway overrepresentation analyses in 1,4-DX-exposed mice. Ingenuity pathway analyses were conducted on the 65 genes that were differentially-expressed in the livers of mice exposed to 5,000 mg/L 1,4-DX in drinking water for four weeks. Pathways shown to be enriched included those involved in DNA damage response (p=8.1E-7) and repair (p=1.0E-2), and hepatocellular carcinoma (p=2.2E-4). Molecules in red have increased expression in 1,4-DX-exposed mice, whereas green molecules are decreased. Blue dashed lines indicate a predicted inhibitory effect, whereas orange dashed lines indicate predicted activation. Grey lines mean that the effect cannot be predicted. The darker the color, the greater the fold change in DEGs between 1,4-DX-exposed and control mice. *PHLDA3*; Pleckstrin homology-like domain family A member 3, *DDIAS*; DNA damage-induced apoptosis suppressor, *GSTP1*; glutathione S-transferase pi 1, *PPPLR10*; protein phosphatase 1 regulatory subunit 10, *BTG2*; BTG anti-proliferation factor, *ID1*; inhibitor of DNA binding 1, *AOX1*; aldehyde oxidase 1, *CCNG1*; cyclin G1, *UGT1A1*; UDP glucuronosyltransferase family 1 member A1.

### 3.4 Untargeted metabolomic analyses

Liver, kidney, urine and feces samples were analyzed by untargeted metabolomics to determine the influence of one- or four-week exposure to 5,000 mg/L 1,4-DX in drinking water on the metabolome. PCA scores plots showed no clear separation between samples from 1,4-DX-exposed mice and control mice **(Supplementary Figure S2).** Univariate volcano plots did not reveal any significantly dysregulated putative metabolites after FDR correction for multiple comparisons; −log10 adjusted p-value threshold cutoff set to 1.3 (i.e., absolute value of 0.05) **(Supplementary Figure S3**).

### 3.5 Targeted metabolomics: quantification of the bile acid biosynthesis pathway

Pathway analysis of liver transcriptomic data of mice exposed to 5,000 mg/L 1,4-DX for four weeks revealed two pathways related to bile acid synthesis, i.e., LPS/IL-1 mediated inhibition of RXR function, and PXR/RXR activation. Furthermore, alterations to bile acid levels are a known biomarker of liver dysfunction. Therefore, we attempted to quantify bile acids in liver tissue and feces from mice exposed to 5,000 mg/L 1,4-DX in drinking water for one or four weeks in order to determine if they were altered in response to 1,4-DX exposure. It was possible to detect 13 bile acids in liver samples and nine bile acids in feces samples (**Figure 5**). Both primary and secondary bile acids were measured, as well as their taurine and glycine amino acid conjugates; however, 1,4-DX exposure did not show any significant difference (p>0.05) in the levels of any of the bile acids relative to control mice (**Figure 5**).

**Figure 5:**
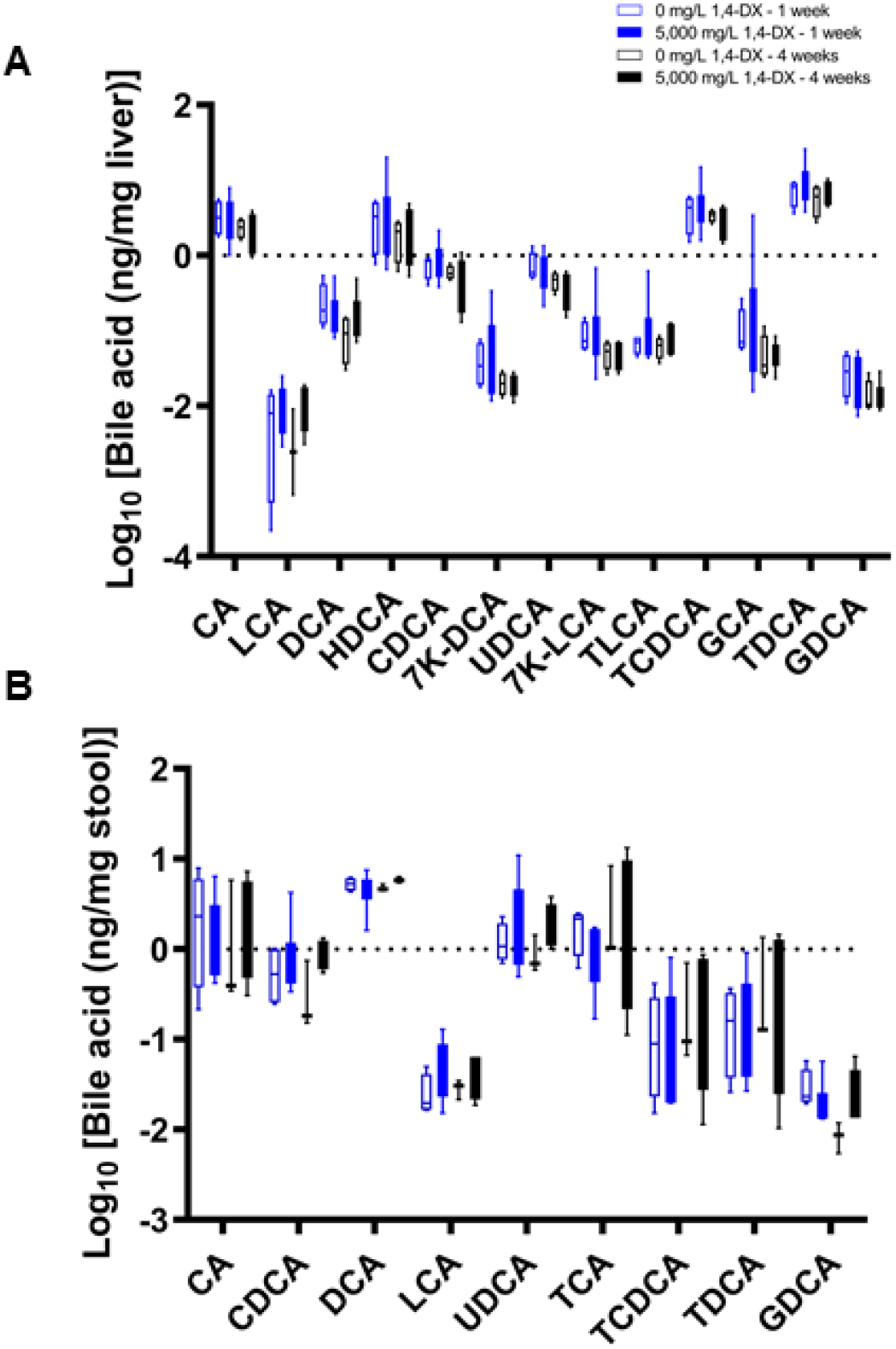
Bile acid concentrations in stool and liver samples from mice exposed to 1,4-DX. Bile acids were quantified by mass spectrometry in liver (**A**) and stool (**B**) samples obtained from mice exposed to 0 mg/L (control) or 5,000 mg/L 1,4-DX in drinking water for one or four weeks. Data are presented as box-and-whisker plots from n=4-6 mice/group. (See Table 1 for bile acid abbreviations.)

## Discussion

Currently, there has been no data showing the mechanism by which 1,4-DX induces liver tumors in experimental animals (Dourson *et al.*, 2017; EPA, 2013). The results of the present study revealed that liver damage starts after one week of exposure to 5,000 mg/L 1,4-DX in mice and is associated with hepatic genetic alterations that may impact DNA damage and hepatocellular carcinoma pathways.

Immunohistochemical analysis of liver tissues using H2AXγ, a marker of DNA double strand breaks, revealed a small but significant increase in the number of hepatocytes with DNA damage in mice exposed to 5,000 mg/L (but not 500 mg/L) 1,4-DX in drinking water for one and four weeks. These results are consistent with previously published research which have shown that 1,4-DX causes hepatic DNA damage in female Sprague-Dawley rats treated with 2,550 and 4,200 mg/kg 1,4-DX, 21 h and 4 h before sacrifice (EPA, 2013; IARC, 1993; Kitchin and Brown, 1990). In addition, 5,000 mg/L exposure to 1,4-DX for one week caused an increased in H2AXγ expression in non-hepatocytes; no such increase in H2AXγ occurred in mice exposed for four weeks. This could indicate that the non-hepatocytes altered their metabolism to account for this chemical stress, or that more non-hepatocytes were recruited from the bloodstream.

CK-7 immunostaining for cellular proliferation revealed increases in precholangiocytes in liver tissue from mice exposed to 5,000 mg/L 1,4-DX for four weeks, but no changes in mice exposed to 500 mg/L for one week. DNA damage was increased in the hepatocytes of mice exposed to 5,000 mg/L 1,4-DX for four weeks (as seen by increased H2AXγ). In addition, an increase in precholangiocytes proliferation was also seen at this exposure concentration. These findings indicate both DNA damage and repair are occurring simultaneously. These are in agreement with the previously-documented changes in the relative mRNA expression of genes involved in cell proliferation after exposure of rats to 5,000 mg/L 1,4-DX for 16 weeks(Gi *et al.*, 2018). No increase in H2AXγ occurred in the kidney tissues of 1,4-DX exposed mice at any dose or time-point, indicating that DNA damage did not occur in the kidney. This contrasts with previous studies that have shown 1,4-DX could elicit kidney toxicity. Long-term oral exposure to higher concentrations of 1,4-DX (10,000 or 25,000 ppm) has been shown to lead to changes in the proximal tubule of kidneys in the rat (Kano, Umeda, Saito, Senoh, Ohbayashi, Aiso, Yamazaki, Nagano and Fukushima, 2008). Other studies have shown degenerative changes to kidney tissues after 1,4-DX exposure, including vacuolar degeneration, and/or focal tubular epithelial regeneration in proximal cortical tubules (NCI, 1978).

Pathway analysis of the differentially-expressed genes from liver samples of mice exposed to 5,000 mg/L 1,4-DX for four weeks revealed that xenobiotic metabolic pathways were the most enriched pathways; these include glutathione-mediated detoxification, NRF2-mediated oxidative stress response and glutathione redox reactions. The observed down-regulation of *Cyp3a16* (human orthologue *CYP3A5*) may have important pathophysiological ramifications. CYP3A is a member of the cytochrome P450 monooxygenase superfamily and is responsible for the oxidative metabolism of numerous drugs. Several studies have shown that inflammation and infection cause suppression of cytochrome P450 levels in various species, including human, rat and mouse (Morgan, 1997; Okamura *et al.*, 2019). In the present study, the down-regulation of *Cyp3a16* may be caused by 1,4-DX-induced inflammation. In support of this notion, our pathway analysis suggested alterations in the “LPS/IL-1-mediated inhibition of RXR function” and “PXR/RXR activation” pathways. Lipopolysaccharide (LPS) induces down-regulation of CYP3A in mice (Li-Masters and Morgan, 2001; Morgan, 2001), and has been associated with reductions in PXR mRNA and protein levels (Beigneux *et al.*, 2002; Xu *et al.*, 2004). In the present study, two out of the four identified genes in the PXR/RXR pathway were down-regulated, and the LPS/IL-1-mediated inhibition of RXR function pathway showed *CYP3A5, SULT2C2, SULT3A1*, and *SULT3A2* to be down-regulated. Glutathione S-transferase genes were up-regulated. These changes are consistent with an oxidative stress response. 1,4-DX exposure also caused an alteration in the melatonin degradation pathway. Melatonin has been shown to influence the immune system and is an endogenous substance that exhibits antioxidative, anti-inflammatory and anti-apoptotic properties (Cichoż-Lach and Michalak, 2014). IPA analysis of the differentially-expressed genes was used to identify the biological functions most affected by 1,4-DX exposure. These analyses revealed DNA damage response and DNA repair functions, and hepatocellular carcinoma. These findings are consistent with our histopathology results which showed increased DNA double strand breaks in the hepatocytes after both one and four weeks of 5,000 mg/L 1,4-DX exposure. *PHLDA3* and *DDAIS* were both enriched within the “DNA damage response and repair functions”. The role of these genes in relation to hepatocellular carcinoma have been previously investigated (Han *et al.*, 2016; Zhang *et al.*, 2015). Specifically, PHLDA3 appears to play a key role in hepatocyte death activated by endoplasmic reticulum stress; it acts as an effector for the execution of apoptosis and facilitates liver injury by inhibiting Akt signaling (Han, Lim, Koo, Kim and Kim, 2016). Overexpression of DDIAS promotes cellular proliferation, colony formation, cellular migration, and *in vivo* tumorigenicity in hepatocellular carcinoma cells (Han, Lim, Koo, Kim and Kim, 2016). DDIAS knockdown attenuates these effects (Zhang, Huang, Wang, Cai and Han, 2015). These mechanisms could also be contributing to the hepatic carcinogenic potential of 1,4-DX.

The untargeted metabolomic analyses showed no significant changes in the metabolome of liver, kidney, urine and feces after 1,4-DX exposure. This is in contrast to a recent publication which reported changes in metabolites and pathways in kidney tissues as early as three weeks after the initiation of exposure to 500 mg/L 1,4-DX in drinking water (Qiu *et al.*, 2019). However, there are no other publications reporting metabolomic data and very limited information is available on the metabolic response to 1,4-DX treatment. Our analysis of differentially-expressed genes indicated that DNA damage and repair functions were activated in the liver, a result supported by our histopathology data. Our results would suggest that that 1,4-DX exposure caused DNA damage in the liver but did not cause cellular death. In addition, we observed evidence of a repair mechanism occurring in the liver of these same mice, i.e., an increased number of precholangiocytes (observed through CK-7 staining).

The DNA damage and repair mechanisms observed in the present study would have been expected to cause changes in the liver metabolome. Similarly, alterations in bile acid levels (which are a hallmark of liver dysfunction) would have been anticipated in the livers of 1,4-DX-treated mice. As such, the lack of 1,4-DX-induced alterations to the metabolome in either untargeted analyses or in targeted analyses of bile acids is enigmatic. It is conceivable that the hepatocellular DNA damage observed by IHC did not affect the processing of bile acids in the liver. Further experiments in mice exposed to 1,4-DX for longer periods of time may reveal how metabolomic changes relate to hepatic damage. It is possible that more significant cellular changes occurring as a consequence of the DNA damage and repair are necessary before metabolomic analyses can detect the biochemical consequences of such alterations.

In the present study, only exposure to the highest concentration of 1,4-DX (i.e., 5,000 mg/L in drinking water) and for the longest duration (i.e., four weeks) caused histological manifestations of DNA damage and repair, increases in the number of hepatocytes showing DNA damage, and increases in the number of precholangiocytes (suggestive of repair). These increases, while significant, were small, suggesting that this exposure level only caused minor hepatic toxicity. Hepatic transcriptome revealed significant genetic changes in the livers of these mice, and pathway analysis of differentially-expressed genes revealed DNA damage and repair metabolic pathways. Genes involved in xenobiotic metabolism signaling were also altered, as were those linked to oxidative stress and melatonin metabolism. Changes in these pathways would be predicted to make the mice vulnerable to liver cancer. An enigmatic observation was that the metabolomes of liver, kidney, feces and urine samples were unaltered in the same groups of mice in which 1,4-DX exposure had caused histological and genetic changes. Our studies reveal the capacity of 1,4-DX to selectively induce damage in the liver. It should be recognized that these studies were of a relatively short duration compared to the exposure that may occur in humans *via* contaminated drinking water.

## Acknowledgements

We would like to thank Dennis Petersen, University of Colorado, Aurora, CO for providing the 4-HNE antibody. We would like to thank Yale School of Public Health for providing financial support for this study.

