## Supplementary File for "Novel insights into the mode of action of 1,4-dioxane using a systems screening approach"

**Supplementary Figure S1**

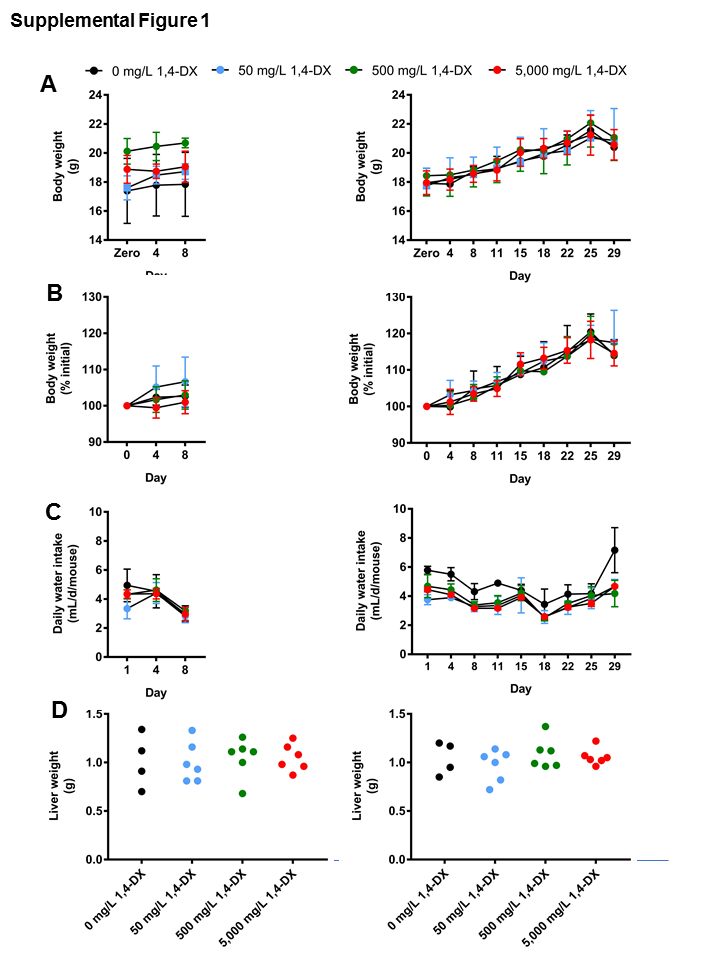

**Supplementary Figure S1**: *Effect of 1,4-DX exposure on body weight, water intake and liver tissue weight.* Mice were exposed to various concentrations of 1,4-DX (0, 50, 500 or 5,000 mg/L) for one (left panel) or four (right panel) weeks. **A, B)** Changes in mouse body weight. Mouse weights were expressed as a percentage of the body weight prior to 1,4-DX exposure. **C)** Daily consumption of water. **D)** Liver weights of mice at the end of the 1,4-DX exposure period. Data are presented as the mean ± standard deviation for each exposure group (n=4-6 mice/group). Differences between the groups at each time-point were examined using one-way ANOVA with the Tukey's *post-hoc* correction for multiple testing. The analysis showed no significant (p>0.05) differences between the groups.

**Supplementary Figure S2**

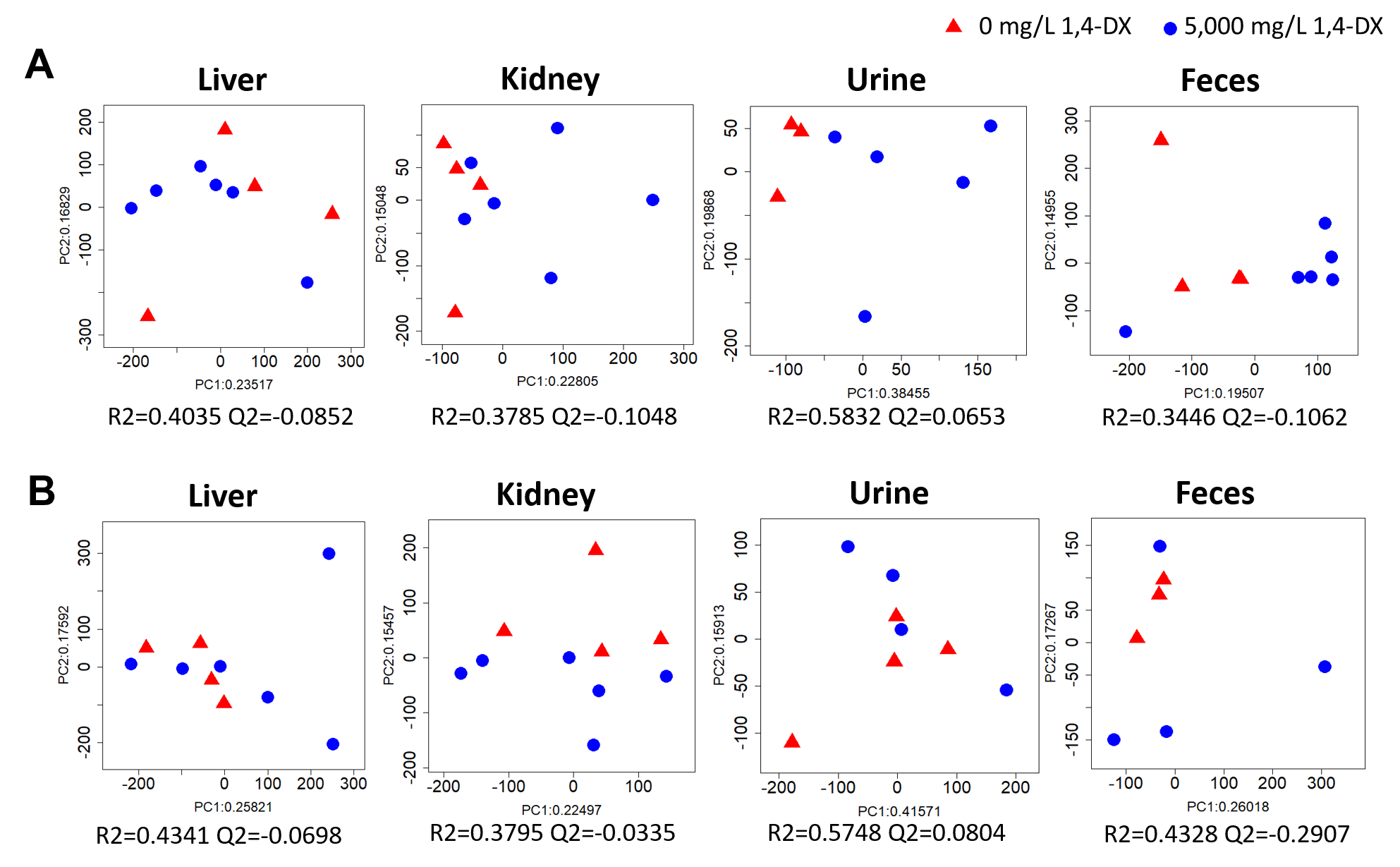

**Supplementary Figure S2:** *Principal components analysis (PCA) of untargeted metabolomics data from mice exposed to 5,000 mg/L 1,4-dioxane (1,4-DX).* PCA scores plots are shown from analyses of liver, kidney, urine, and feces samples obtained from mice exposed to 0 or 5,000 mg/L 1,4-DX in drinking water for one (**A**) or four **(B**) weeks. Liver and fecal samples were analyzed by HILIC-MS and RPLC-MS in both positive and negative electrospray ionization (ESI) modes, and data were combined for PCA. Kidney and urine samples were analyzed by RPLC-MS positive ESI mode. Model statistics are shown below each plot where R2 represents the percentage of explained variance, and Q2 represents the percentage of predicted variation.

**Supplementary Figure S3**

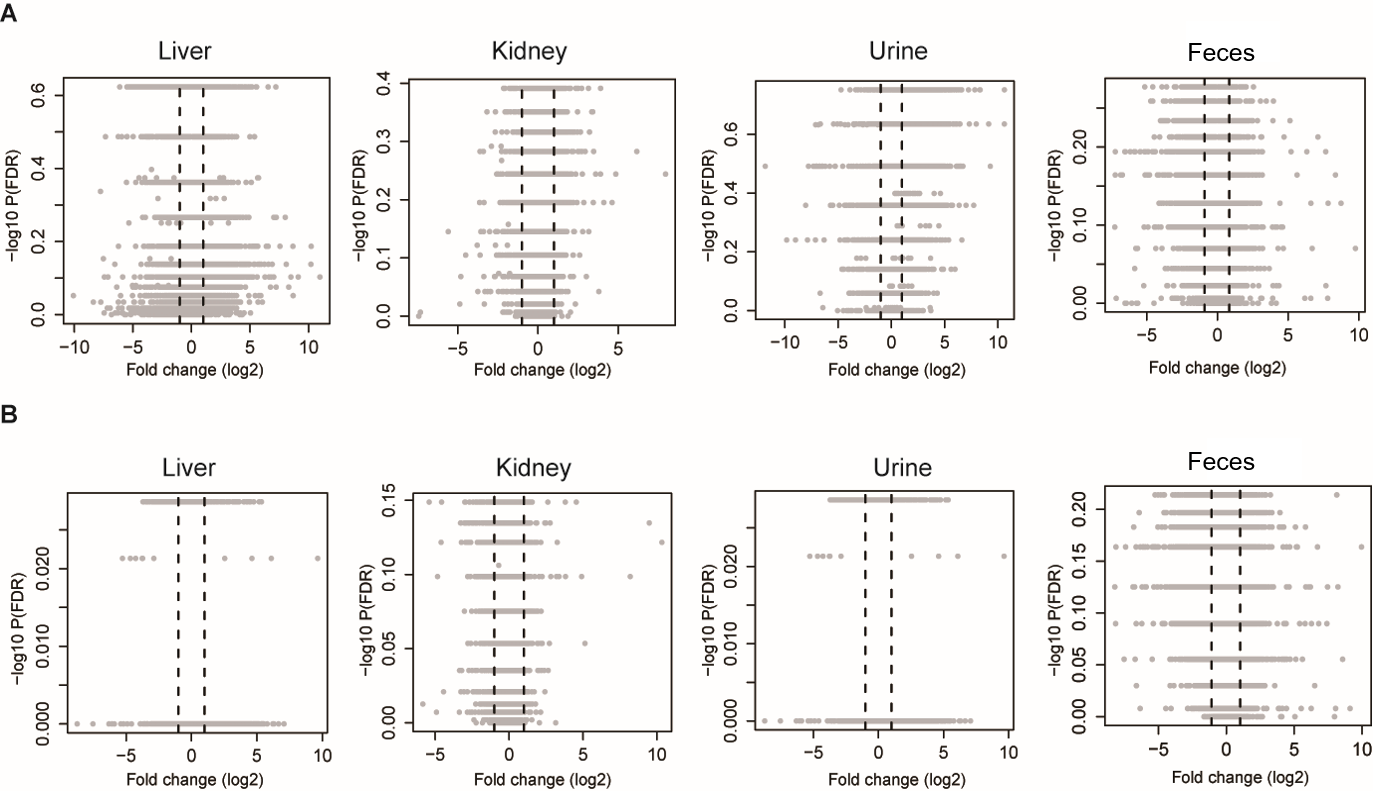

**Supplementary Figure S3:** *Volcano plots of metabolomics data from mice exposed to 5,000 mg/L 1,4-dioxane (1,4-DX) for one (****A****) or four (****B****) weeks.*Liver and feces samples were analyzed by HILIC-MS and RPLC-MS in both positive and negative electrospray ionization (ESI) modes), and data were combined for Volcano plot development. Kidney and urine samples were analyzed by RPLC-MS positive ESI mode. Vertical dashed lines show up-regulated (positive values) or down-regulated (negative values) with log_2_ fold change at >1 or <-1.

**SupplementaryTable 1**: *Influence of 1,4-dioxane exposure (1,4-DX) on body weight, absolute and relative liver weight, and 1,4-dioxane (1,4-DX) consumption.*

| 1,4-DX  (mg/L) ^a^ | Exposure duration  (wk) | No. of mice | Body weight ^b^  (g) | Liver weight  (g) | Liver weight /Body weight ^c^ (%) | Water consumption  (ml/mouse/d) | 1,4-DX intake | |
| --- | --- | --- | --- | --- | --- | --- | --- | --- |
|  |  |  |  |  |  |  | Daily rate ^d^  (µg/g) | Total ^e^  (µg) |
| 0 | 1 | 4 | 17.8±2.2 | 1.0±0.3 | 5.7±1.2 | 4.1±0.8 | 0 | 0 |
| 50 | 1 | 6 | 18.7±0.5 | 1.0±0.2 | 5.3±1.0 | 3.5±0.1 | 9.6 ± 0.4 | 67.5 ± 2.8 |
| 500 | 1 | 6 | 20.7 ±0.3 | 1.1±0.2 | 5.1±0.9 | 4.0±0.3 | 99 ± 8 | 693 ± 58 |
| 5,000 | 1 | 6 | 19.1±1.1 | 1.1±0.1 | 5.5±0.5 | 3.9±0.1 | 1030 ±38 | 7,210 ± 270 |
| 0 | 4 | 4 | 20.4±0.6 | 1.0±0.2 | 5.1±0.7 | 4.9±0.1 | 0 | 0 |
| 50 | 4 | 6 | 20.9±2.2 | 1.0±0.2 | 4.6±0.4 | 3.6±0.4 | 9.3 ± 0.5 | 261 ± 14 |
| 500 | 4 | 6 | 21.1±1.6 | 1.1±0.2 | 5.2±0.6 | 3.8±0.4 | 96 ± 13 | 2,690 ± 360 |
| 5,000 | 4 | 6 | 20.6±1.0 | 1.1±0.1 | 5.1±0.2 | 3.6±0.0 | 927 ± 35 | 25,950 ± 980 |

^a^	1,4-DX concentration in drinking water.

^b^	Weight of mouse immediately prior to being euthanized, i.e., terminal body weight.

^c^	Ratio of liver weight to body weight, expressed as a percentage.

^d^	Daily 1,4-DX intake rate = (concentration of 1,4-DX in drinking water (µg/mL)) x (average water consumption (mL/d)) x (exposure frequency (d)) / (body weight (g) x exposure duration (d)).

^e^	Total 1,4-DX consumed = (daily 1,4-DX intake rate) x (exposure duration (d)).

Data are presented as the mean ± standard deviation.
